# Internal structure of *Mycoplasma mobile* gliding machinery analyzed by negative staining electron tomography

**DOI:** 10.1101/2024.02.08.578511

**Authors:** Minoru Fukushima, Takuma Toyonaga, Yuhei O. Tahara, Daisuke Nakane, Makoto Miyata

## Abstract

*Mycoplasma mobile* is a parasitic bacterium that forms gliding machinery on the cell pole and glides on a solid surface in the direction of the cell pole. The gliding machinery consists of both internal and surface structures. The internal structure is divided into a bell at the front and chain structure extending from the bell. In this study, the internal structures prepared under several conditions were analyzed using negative-staining electron microscopy and electron tomography. The chains were constructed by linked motors containing two complexes similar to ATP synthase. A cylindrical spacer with a maximum diameter of 6 nm and a height of 13 nm, and anonymous linkers with a diameter of 0.9–8.3 nm and length of 5–30 nm were found between motors. The bell is bowl-shaped and features a honeycomb surface with a periodicity of 8.4 nm. The chains of the motor are connected to the rim of the bell through a wedge-shaped structure. These structures may play roles in the assembly and cooperation of gliding machinery units.

**Significance:** *Mycoplasma mobile*, a parasitic bacterium, glides on solid surfaces at speeds of up to 4.0 μm per second through a specialized mechanism. The gliding machinery, located at one pole of the cell, is composed of surface legs and internal motor structures. The force generation unit within the internal structure evolved from ATP synthase. This study aimed to clarify the entire architecture of the gliding machinery using electron tomography.

## Introduction

*Mycoplasma mobile* is a parasitic bacterium found on the gills of freshwater fish [1,2], forming a gliding machinery on the cell pole and glides on the solid surface in the direction of the cell pole at a maximum speed of 4 μm/s (Fig. 1). This gliding mechanism, observed only in *M. mobile* and its related species, repeatedly grabs and pulls sialylated oligosaccharides [3-6] onto the host cell surface via ATP hydrolysis [7-10]. Although this mechanism appears similar to that of *Mycoplasma pneumoniae*, they do not share the proteins required for gliding [2,11-18]. The gliding machinery of *M. mobile* and its related species consists of internal and surface structures [19-23]. The main protein components of the surface structures are Gli123, Gli349, Gli521, and Gli42 [10,24-27]. Gli123, Gli349, and Gli521 are involved in complex assembly, binding to sialylated oligosaccharides, and transmitting gliding forces, respectively. The internal structure consists of a bell, which is a rigid structure that exists in front of the cell, and chains of twin motors that evolved from F-type ATP synthase. The structures of Gli349, Gli521, and twin motors have been analyzed; however, the structure of the bell and the entire internal structure remain unclear [19,21,22], which are necessary to understand the gliding mechanism. In this study, the internal structures prepared under several conditions were analyzed using negative staining and electron tomography.

**Fig. 1.**
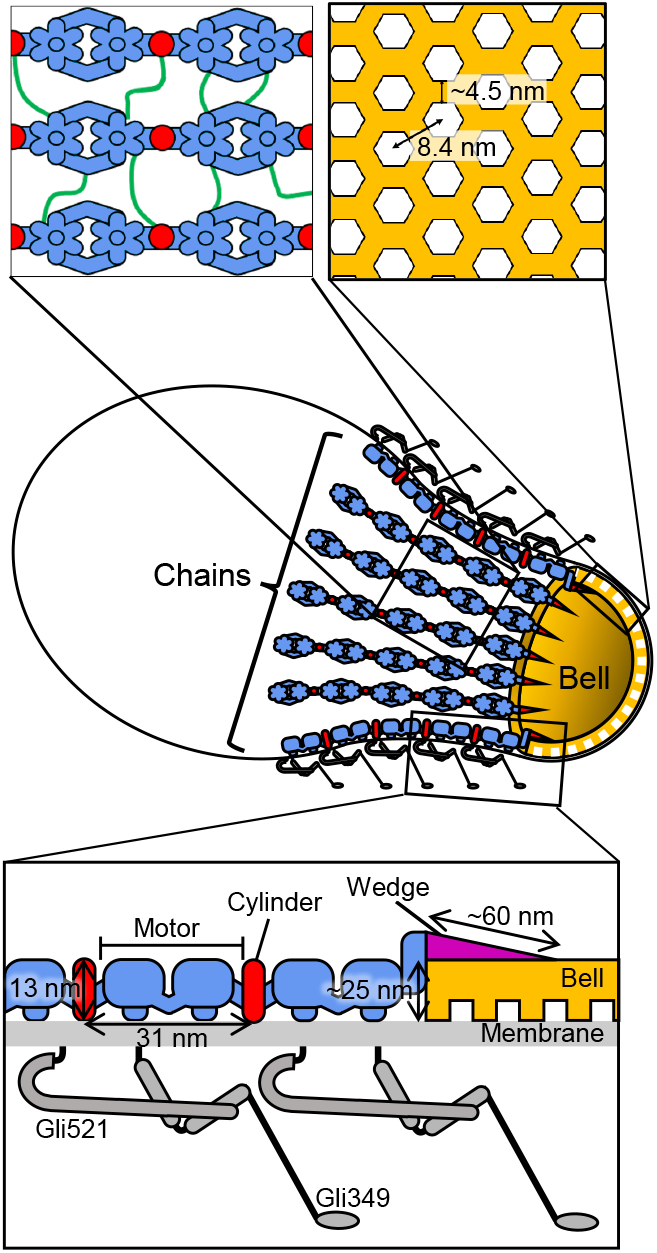
Schematic of *Mycoplasma mobile* gliding machinery including results of this study. A whole-cell approximately 0.8 μm long is presented in the middle. Internal part of machinery consists of bowl-shaped bell featuring honeycomb surface (yellow) and 46 chains with 17 particles each and 430 nm long. Only five motors are shown for each chain. Magnified schematics of chain and bell are shown in upper left and right panels, respectively. Anonymous linkers and cylinders are colored green and red, respectively. A magnified schematic of surface section is shown in bottom panel. Surface structure contains two large proteins, Gli521 and Gli349, which act as a crank for force transmission and a receptor of sialylated oligosaccharide, respectively.

## Materials and Methods

### Bacterial culture

*M. mobile* 163K (ATCC 43663) with an OD_600_ at 0.08 was inoculated to nine-fold volume of fresh Aluotto medium and grown at 25°C for two days [28,29].

### Sample preparation

Cultured cells were collected by centrifugation at 10,000 × g at 25°C for 5 min and suspended in Aluotto medium for a 30-fold concentration of the culture [28,29]. A cell suspension of 5 μL was placed on a carbon-coated electron microscope (EM) grid for 10 min. The suspension was removed, and the grids were treated twice with 5 μL of a detergent solution (0.1% (v/v) Triton X-100, 0.1 mg/mL DNase I, 5 mM MgCl_2_ and 5 mM CaCl_2_ in PBS consisting of 75 mM sodium phosphate (pH 7.3), and 68 mM NaCl) for 1 min at 25°C. The grids were then washed with 10 μL PBS and stained with 2% phosphotungstic acid and air-dried. Permeabilized cells were prepared by a single treatment with detergent solution. Chemically fixed cells were prepared by treating twice with the detergent solution including 0.2% (v/v) Triton X-100, and fixed with 2% (v/v) glutaraldehyde for 5 min at 25°C.

### Image acquirement and analysis

The samples were observed using a Talos F200C G2 transmission electron microscope (Thermo Fisher Scientific, Waltham, MA, USA) at 200 kV with a 4k × 4k Ceta 16M CMOS detector (Thermo Fisher Scientific). Single-axis tilt series were collected covering an angular range from -65° to +65° or -60° to +60° at 6- to 8-μm underfocus using the Tomography software package (Thermo Fisher Scientific). Subtomogram averaging was done the eman 2.99 software package. Tomographic reconstruction was performed using IMOD 4.11.8 software package. The fast Fourier transform (FFT) was calculated using ImageJ v1.53g. Finally, 3D-rendering and segmentation processes were performed using Chimera 1.15 software package.

## Results

### Visualization of isolated gliding machinery

Cultured *M. mobile* cells that adhered to an EM grid were extracted using a detergent and subsequently stained. This process removes the cell membrane and cytoplasm, thereby exposing the internal structures. Bell and chain parts were observed (Fig. 2A–C). Tilt images were captured in increments of either 1°or 3°, reaching 60° or 65° (Movies 1 and 2). The tomograms were then reconstructed from these tilt series (Fig. 2D–F). The features evident in the 2D images are consistent with those reported previously [20,22,23]. The reconstruction revealed an internal structure with a maximum thickness of 50.4 nm in the desiccated state (Fig. 2G).

**Fig. 2.**
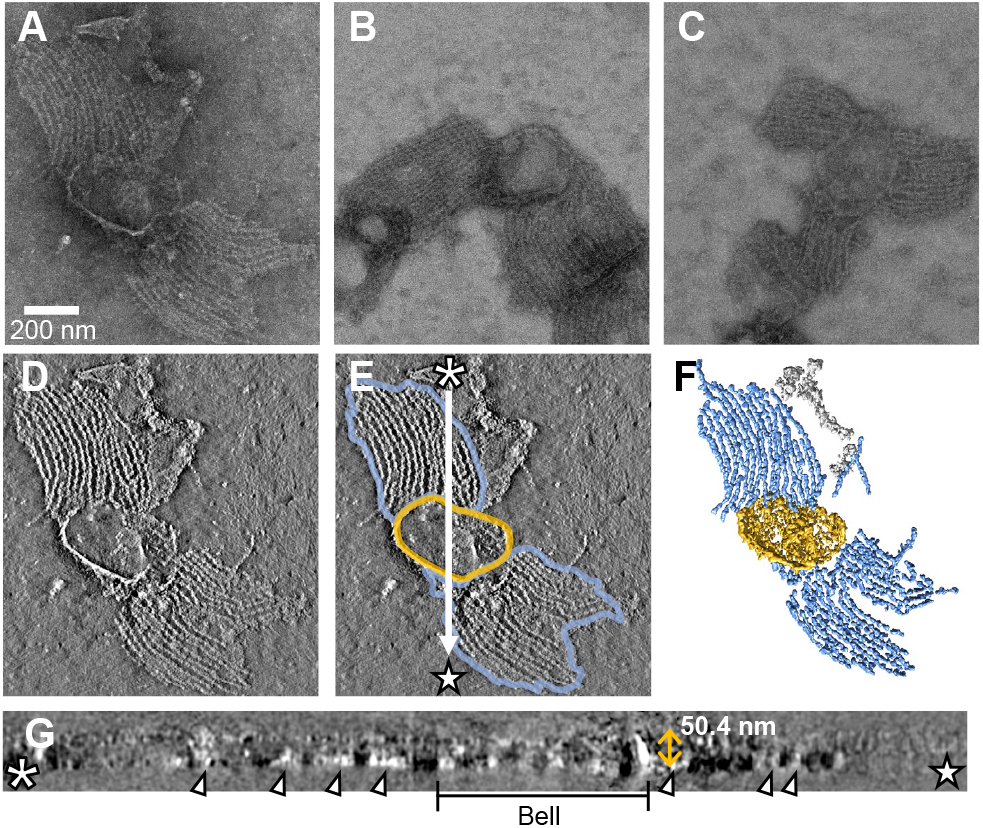
Electron tomogram of internal machinery exposed by Triton X-100 treatment. (A–C) Single image from individual tilt series. (D) A 1.5 nm-thick sliced image of tomogram shown in (A). (E) Schematic image overlaid on image shown in (D). Regions surrounded by blue and yellow lines indicate chain bundles and bell, respectively. (F) 3D-rendered image of tomogram shown in (D). (G) Cross section along white arrow in (E). Bell 50.4 nm high and chains are marked by a black bar and open triangles, respectively.

### Bell structure

The bell is known to have a honeycomb structure; however, its overall architecture remains unclear [23]. Thus, we focused on the bells that had been exposed to two rounds of detergent treatment on an EM grid (Fig. 3). Tilt series of the exposed bells were acquired across a range of -65° to +65° and subsequently reconstructed into tomograms. The resulting reconstructed bell has a thickness of approximately 50 nm. The FFT of the horizontal image revealed three distinct directional periodicities (Fig. 3H). Each periodicity is oriented at a 60-degree interval. The periodicity in all three directions was 8.4 nm (Figs. 1 and 3H).

**Fig. 3.**
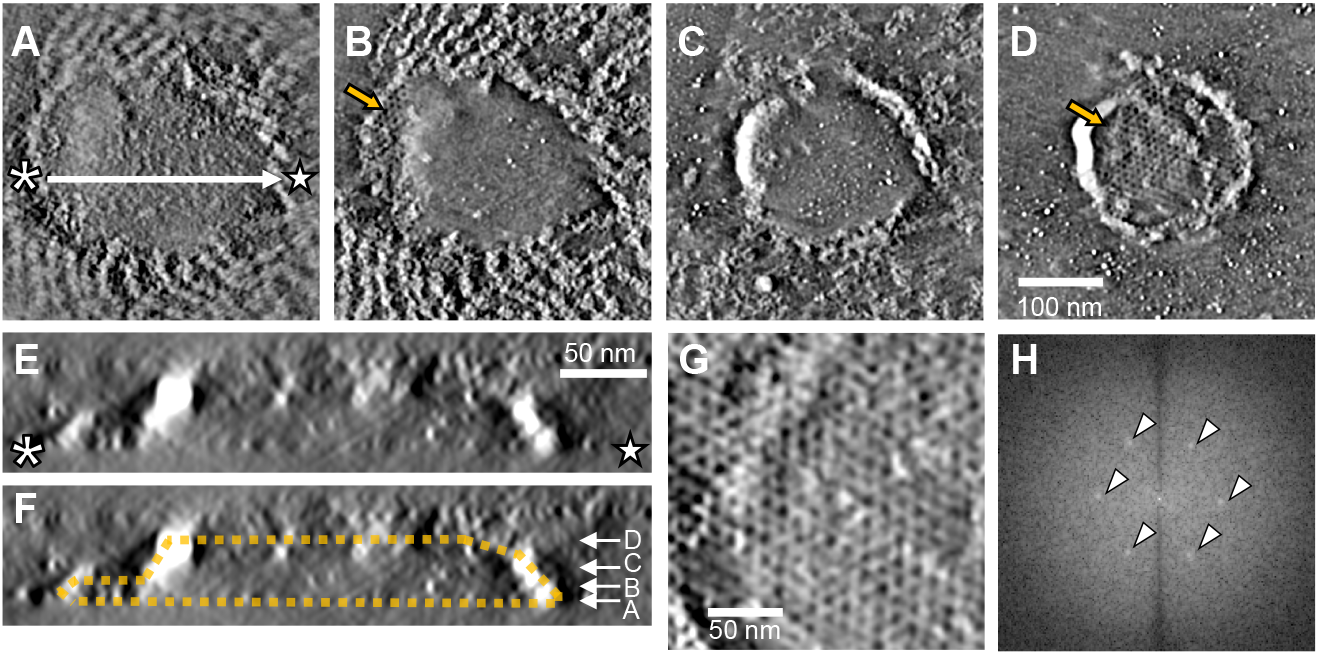
Honeycomb structure of bell. (A–D) Slice images of bell at different heights. Honeycomb structures are observed at surface of bell indicated with yellow arrow. (E) Cross section along position of white arrow line in (A). (F) Marked image of (D). Bell is marked by a yellow polyhedral. Height positions of panels (A–D) are indicated with white arrows. (G) Surface image of bell. (H) Fast Fourier transform image of (G) processed with Tukey window function, which shows six dots with a periodicity of 8.42 nm marked by open triangles.

Determining whether the holes within the honeycomb structure permeated through the entire bell height or were limited to its surface remained challenging, owing to the inadequately adjustable staining conditions (Fig. 3A–D). The bell was deformed due to drying during sample preparation. To observe the bells under more natural conditions, we applied glutaraldehyde fixation before drying (Fig. 4A–C). This approach visualizes the rim of a bowl-shaped bell (Fig. 1). The chains and bell rims were characterized by distinct densities (Fig. 4D). The bells exhibited bowl structures with some distortions (Fig. 4E and F). A chemically fixed bell has a height of approximately 100 nm and a major axis of approximately 490 nm. In this cell, 46 chains of 432 ± 76 nm (n = 153) extend radially outward from the bell at intervals of approximately 24 nm.

**Fig. 4.**
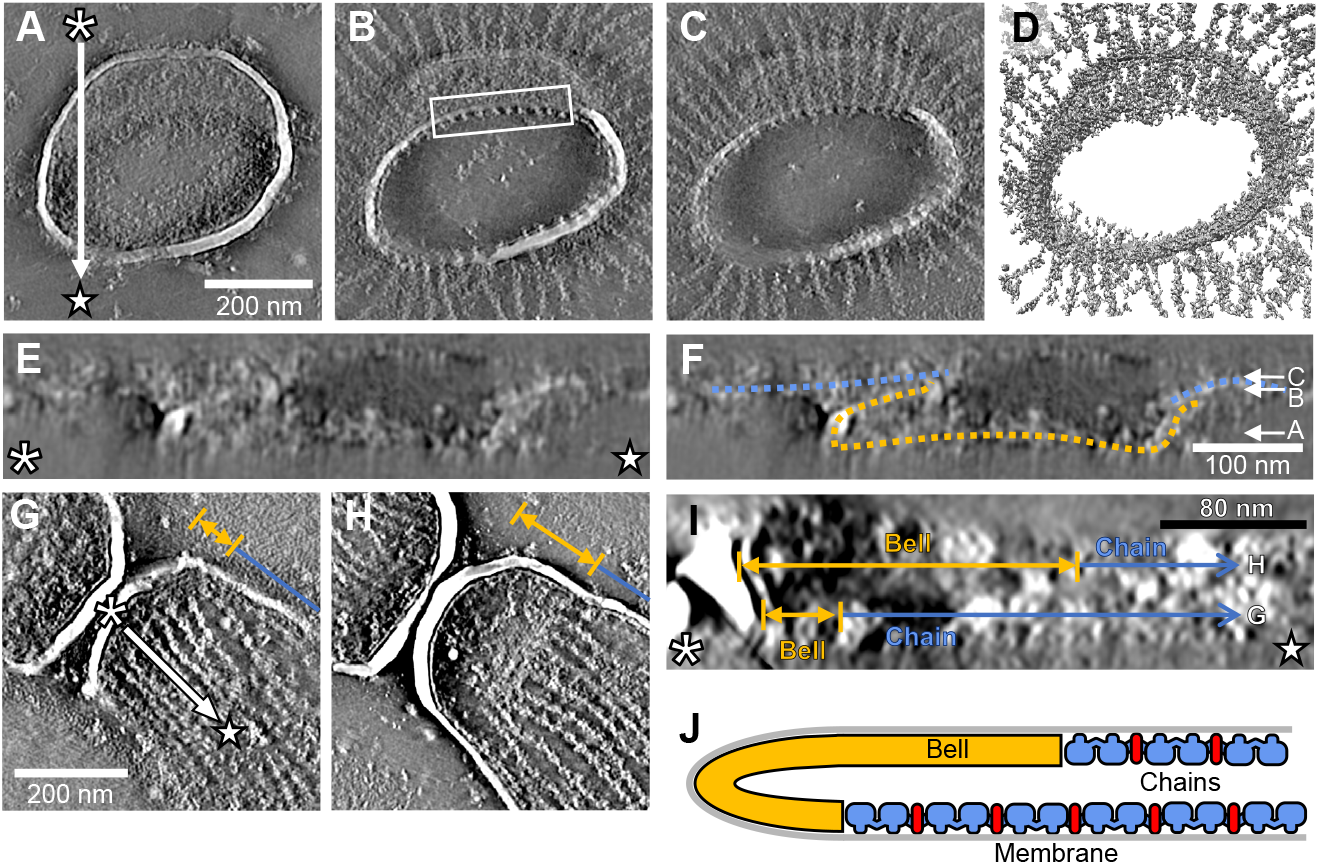
Bell as a bowl. (A–C) Slice images of tomogram from chemically fixed internal machinery. Bell is featured by circular rim. (D) 3D-rendered image of tomogram shown in (A). (E) Cross section along white arrow in (A). (F) Marked image of (E). Bell is outlined with yellow line. Height positions of panels (A–D) are indicated with white arrows. (G and H) Slice images of permeabilized cell tomogram. Two layers of bell and chains are shown. Axial positions of bell and chains indicated with yellow arrows and blue lines, respectively. (I) Cross section along white arrow in tomogram (G). Height positions of slices (G) and (H) indicated with yellow and blue lines. (J) Schematic of permeabilized cell suggested from images (G–I).

Additionally, to gain insight into the structure of the intracellular gliding machinery, we analyzed specimens that preserved some cell membranes with reduced detergent extraction. (Fig. 4G and H). The focused cell was approximately 80 nm high, and the bowl was better retained than in the fully extracted specimens (Fig. 4I and J). The chains extend from the bell in a uniform direction, with a distinct orientation from the fully extracted state.

### Chain structure

We focused on chain structures. Small densities were observed between the two particles, *i*.*e*. twin motors (Fig. 5A). Particle images were obtained from four tomograms. Each particle contained two twin motors that formed a chain. A structure with a resolution of 31.0 Å (Fourier shell correlation [FSC] = 0.143) was obtained through subtomogram averaging of 718 particles (Fig. 5B). Ribbon models of the F_1_-ATPase from *Bacillus* sp. PS3 (PDB ID 7XKQ) fitted well into the reconstructed chain model, indicating that the negatively stained chain maintained its original structure (Fig. 5C). Several novel features were found here (Fig. 1). A cylindrical structure with a height of approximately 13 nm and a maximum diameter of 6 nm was observed at the connecting part of the twin motors. The amorphous filaments were visualized as structures linking neighboring chains. Each filament had a length of 5–30 nm and width of 0.9–8.3 nm. Multiple filaments are extended from a single twin motor (Fig. 5D). Subtomogram averaging of the single-unit twin motors was performed using 1341 particles collected from five tomograms. Averaged structure with a resolution of 24.3 Å (FSC = 0.143) revealed several protrusions extending from the side of F_1_-like hexamers, as previously reported (Fig. 5E) [19].

**Fig. 5.**
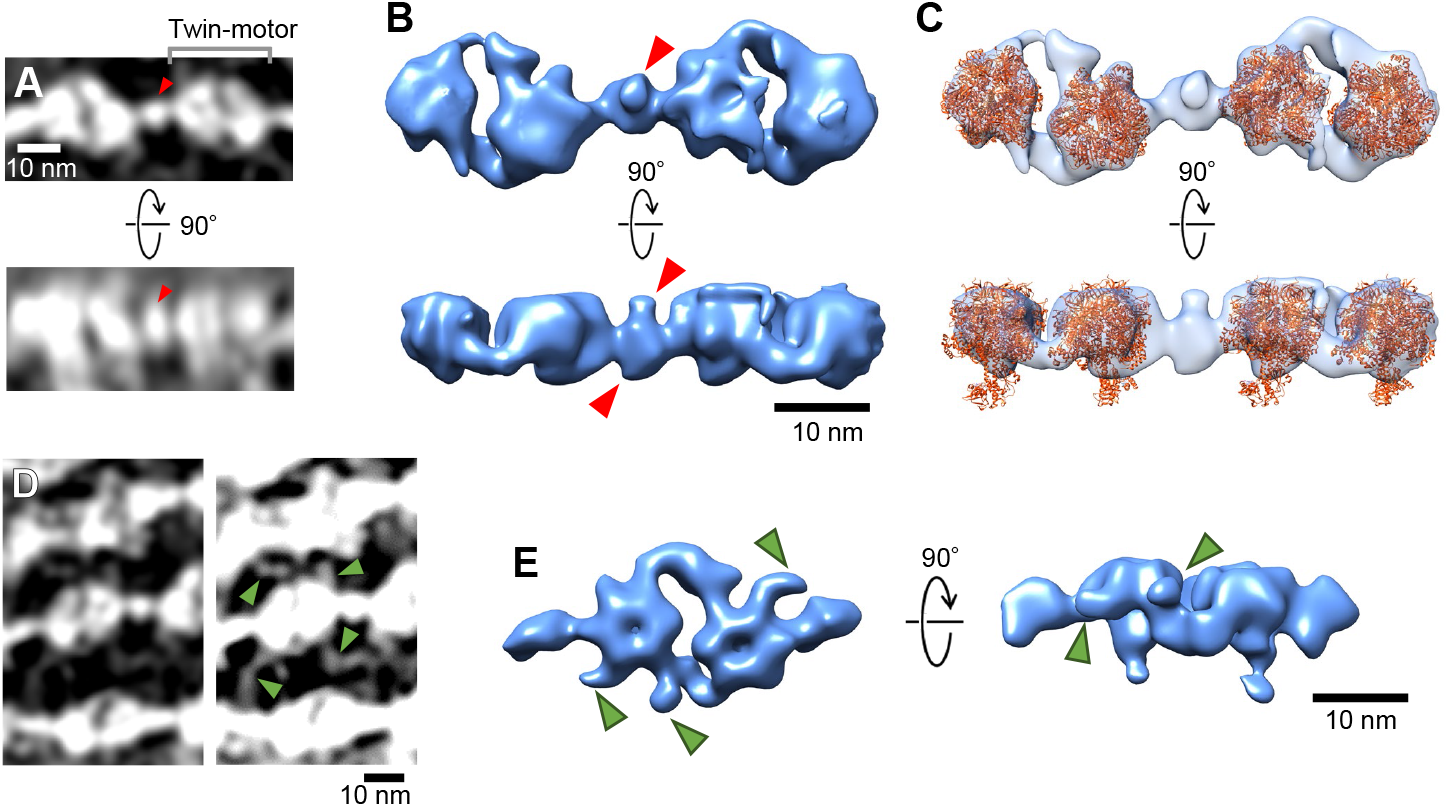
Reconstructed chains. (A) Chain slices focusing on protrusion between two motors. Lower image shows side view of upper image. Protrusions are indicated with red triangle. (B) 3D model of chain reconstructed by subtomogram averaging of 718 particles. Cylinder indicated with red triangle. (C) Superposition of atomic model of *Bacillus* sp. PS3 F_o_F_1_-ATPase F_1_ domain (PDB ID 7XKQ). (D) Chain slices focusing on filamentous structures. Filaments are indicated with green triangles. (E) 3D model of twin motor reconstructed by subtomogram averaging of 1341 particles. Multiple protrusions are extending from sides of F_1_-like hexamers as indicated with green triangles.

Terminal structures of the chains were observed at the edge of the bell in the chemically fixed specimens (Figs. 1 and 6A). These structures had wedge-like shapes and extended along the inner surface of the bell (Fig. 6B and C). The length of each wedge was distributed 60–67 nm. (Fig. 6D).

**Fig. 6.**
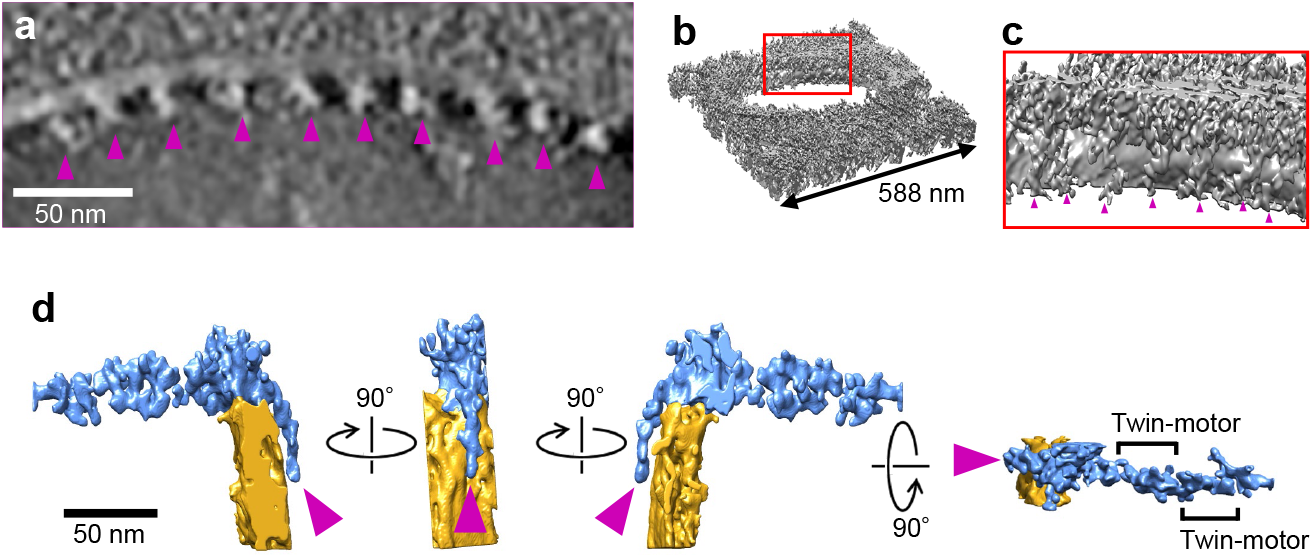
Wedge structure connecting chain to bell. (A) Magnified image of boxed area shown in Fig. 4. Base structures of chain indicated with magenta triangle. Chains are not visible at this height. (B) 3D rendered model of tomogram. (C) Magnified image of boxed area in (B). Wedge structures bind to bell as indicated with magenta triangles. (D) Single unit of structure containing wedge (blue), chain (blue), and part of bell (yellow), viewed from various angles. Wedge is found at a chain terminal as shown in rightmost image indicated with magenta triangle.

## Discussion

### Novel structures and its role

The structure of the gliding machinery is being clarified, focusing on surface proteins such as Gli349 and internal bells and chains [10,19,21,22,24-27]. However, little is known about the structures that assemble this machinery, even though they are essential for clarifying the gliding mechanism. The results obtained in this study may provide clues for elucidating their structures (Fig. 1).

The name “bell” is derived from the first name for the internal structure of the gliding machinery, namely the “jellyfish” structure [23]. The entire bell is shown as a bowl of uniform thickness (Fig. 4). The honeycomb-repeating structure of the bell suggests that it is a small protein, possibly a polymer of MMOB4860 [21,23]. *M. mobile* cells always have one or more gliding machinery, suggesting that the formation of this machinery is tightly coupled with cell proliferation [30,31]. We speculate that the controlled polymerization of bell component proteins is the first step in the formation of *de novo* gliding machinery that couples with cell division.

The twin motors, which are force generation units in the internal structure, are vertically connected to form a chain. In the image reconstructed by cryotomography, these chains were aligned, facing the same plane as that of the membrane [22]. Because the isolated chainbell complex contained no obvious membrane proteins, the chain was expected to bind to an unidentified membrane protein. The cylinders identified in this study may have been linked to this membrane protein. Negative-staining EM images of the isolated internal structure revealed sheets of individual chains that were almost equally spaced, suggesting a lateral linkage of the chains [22,23]. This structure may be the linker that connects the chains identified in this study. We revealed that the bell is a bowl with a honeycomb structure and that the novel wedge anchors the chains to the bell. The number of chains, 46, was larger than that in the previous analysis, which ranged from 15 to 35 [22]. This difference suggests that the number and length of chains may change with the growth conditions.

### Component proteins

Here, we discuss the possible components of the three novel structures identified in this study [19,21,32]. Open reading frames (ORFs) and MMOBs 1620–1670, 0150, 4530, 4860, and 5430 were previously identified as component proteins of isolated internal structures [23]. Excluding the ORFs identified as twin motor component proteins, MMOBs 1640, 1650, 0150, 4860, and 5430 remain unknown [19]. Previous immunofluorescence studies have shown localization of MMOB4860 at the cell front, suggesting that this protein corresponds to the wedge [23]. MMOBs 1640, 1650, and 5430 have been shown to localize to the cell neck based on fluorescent protein fusion results, making them candidates for novel structural components [21].

## Concluding remarks

The results obtained in this study suggest that the bell serves as the starting point for the formation of the gliding machinery and as a structural anchor. When gliding machinery is formed, the component proteins form a honeycomb structure by polymerization, which serves as the starting point of gliding machinery formation, and when in motion, it serves as an anchor for the polarity of the machinery to work cooperatively.

## Author Contributions

MF did experiments. MF, TT, YOT, DN did data analyses. MF and MM wrote a draft. All discussed the story and completed the manuscript.

## Data Availability Statements

The data underlying this article will be deposited to EMDB.

## Acknowledgements

We thank Hideki Nakagawa and Junko Shiomi at Osaka Metropolitan University for their technical help, and the Equipment Sharing Center for Advanced Research and Innovation for the use of the Talos F200C G2 transmission electron microscope and related software.

This study was supported by Grants-in-aid for scientific research (A) (JP17H01544), a JST CREST grant (JPMJCR19S5) to MM, and JST, the establishment of university fellowships for the creation of science technology innovation, Grant Number JPMJFS2138 to MF.

## Movies

Movie 1. Tomography of isolated internal structures. Images every 3° from -60° to +60° are assembled. Horizontal edge of movie is 1540 nm.

Movie 2. Slice movie of isolated internal structure reconstructed from a tilt series. Slice images 75.2 nm high are stacked for 50 slices. Horizontal edge of movie is 1540 nm.

## References

[1] Miyata, M. Unique centipede mechanism of Mycoplasma gliding. Annu Rev Microbiol 64, 519–537 (2010). 10.1146/annurev.micro.112408.134116

[2] Miyata, M., Hamaguchi, T. Prospects for the gliding mechanism of Mycoplasma mobile. Curr Opin Microbiol 29, 15–21 (2016). 10.1016/j.mib.2015.08.010

[3] Hamaguchi, T., Kawakami, M., Furukawa, H., Miyata, M. Identification of novel protein domain for sialyloligosaccharide binding essential to Mycoplasma mobile gliding. FEMS Microbiol Lett 366, fnz016 (2019). 10.1093/femsle/fnz016

[4] Kasai, T., Hamaguchi, T., Miyata, M. Gliding motility of Mycoplasma mobile on uniform oligosaccharides. J Bacteriol 197, 2952–2957 (2015). 10.1128/jb.00335-15

[5] Kasai, T., Nakane, D., Ishida, H., Ando, H., Kiso, M., Miyata, M. Role of binding in Mycoplasma mobile and Mycoplasma pneumoniae gliding analyzed through inhibition by synthesized sialylated compounds. J Bacteriol 195, 429–435 (2013). 10.1128/JB.01141-12

[6] Nagai, R., Miyata, M. Gliding motility of Mycoplasma mobile can occur by repeated binding to N-acetylneuraminyllactose (sialyllactose) fixed on solid surfaces. J Bacteriol 188, 6469–6475 (2006). 10.1128%2FJB.00754-06

[7] Mizutani, M., Tulum, I., Kinosita, Y., Nishizaka, T., Miyata, M. Detailed analyses of stall force generation in Mycoplasma mobile gliding. Biophys J 114, 1411–1419 (2018). 10.1016/j.bpj.2018.01.029

[8] Kinosita, Y., Miyata, M., Nishizaka, T. Linear motor driven-rotary motion of a membrane-permeabilized ghost in Mycoplasma mobile. Sci Rep 8, 11513 (2018). 10.1038/s41598-018-29875-9

[9] Kinosita, Y., Nakane, D., Sugawa, M., Masaike, T., Mizutani, K., Miyata, M., et al. Unitary step of gliding machinery in Mycoplasma mobile. Proc Natl Acad Sci U S A 111, 8601–8606 (2014). 10.1073/pnas.1310355111

[10] Uenoyama, A., Miyata, M. Gliding ghosts of Mycoplasma mobile. Proc Natl Acad Sci USA 102, 12754–12758 (2005). 10.1073/pnas.0506114102

[11] Nakane, D. Rheotaxis in Mycoplasma gliding. Microbiol Immunol 67, 389–395 (2023). 10.1111/1348-0421.13090

[12] Nakane, D., Kabata, Y., Nishizaka, T. Cell shape controls rheotaxis in small parasitic bacteria. PLoS Pathog 18, e1010648 (2022). 10.1371/journal.ppat.1010648

[13] Nakane, D., Murata, K., Kenri, T., Shibayama, K., Nishizaka, T. Molecular ruler of the attachment organelle in Mycoplasma pneumoniae. PLoS Pathog 17, e1009621 (2021). 10.1371/journal.ppat.1009621

[14] Mizutani, M., Sasajima, Y., Miyata, M. Force and stepwise movements of gliding motility in human pathogenic bacterium Mycoplasma pneumoniae. Front Microbiol 12, 747905 (2021). 10.3389/fmicb.2021.747905

[15] Miyata, M., Robinson, R.C., Uyeda, T.Q.P., Fukumori, Y., Fukushima, S.I., Haruta, S., et al. Tree of motility - A proposed history of motility systems in the tree of life. Genes Cells 25, 6–21 (2020). 10.1111/gtc.12737

[16] Mizutani, M., Miyata, M. Behaviors and energy source of Mycoplasma gallisepticum gliding. J Bacteriol 201 (2019). 10.1128/JB.00397-19

[17] Miyata, M., Hamaguchi, T. Integrated information and prospects for gliding mechanism of the pathogenic bacterium Mycoplasma pneumoniae. Front Microbiol 7, 960 (2016). 10.3389/fmicb.2016.00960

[18] Nakane, D., Kenri, T., Matsuo, L., Miyata, M. Systematic structural analyses of attachment organelle in Mycoplasma pneumoniae. PLoS Pathog 11, e1005299 (2015). 10.1371/journal.ppat.1005299

[19] Toyonaga, T., Kato, T., Kawamoto, A., Kodera, N., Hamaguchi, T., Tahara, Y.O., et al. Chained structure of dimeric F_1_-like ATPase in Mycoplasma mobile gliding machinery. mBio 12, e0141421 (2021). 10.1128/mBio.01414-21

[20] Kobayashi, K., Kodera, N., Kasai, T., Tahara, Y.O., Toyonaga, T., Mizutani, M., et al. Movements of Mycoplasma mobile gliding machinery fetected by high-dpeed stomic force microscopy. mBio 12, e0004021 (2021). 10.1128/mBio.00040-21

[21] Tulum, I., Kimura, K., Miyata, M. Identification and sequence analyses of the gliding machinery proteins from Mycoplasma mobile. Sci Rep 10, 3792 (2020). 10.1038/s41598-020-60535-z

[22] Nishikawa, M., Nakane, D., Toyonaga, T., Kawamoto, A., Kato, T., Namba, K., et al. Refined mechanism of Mycoplasma mobile gliding based on structure, ATPase activity, and sialic acid binding of machinery. mBio 10, e02846–02819 (2019). 10.1128/mBio.02846-19

[23] Nakane, D., Miyata, M. Cytoskeletal “jellyfish” structure of Mycoplasma mobile. Proc Natl Acad Sci U S A 104, 19518–19523 (2007). 10.1073/pnas.0704280104

[24] Tulum, I., Yabe, M., Uenoyama, A., Miyata, M. Localization of P42 and F_1_-ATPase alpha-subunit homolog of the gliding machinery in Mycoplasma mobile revealed by newly developed gene manipulation and fluorescent protein tagging. J Bacteriol 196, 1815–1824 (2014). 10.1128/JB.01418-13

[25] Uenoyama, A., Miyata, M. Identification of a 123-kilodalton protein (Gli123) involved in machinery for gliding motility of Mycoplasma mobile. J Bacteriol 187, 5578–5584 (2005). 10.1128%2FJB.187.16.5578-5584.2005

[26] Seto, S., Uenoyama, A., Miyata, M. Identification of a 521-kilodalton protein (Gli521) involved in force generation or force transmission for Mycoplasma mobile gliding. J Bacteriol 187, 3502–3510 (2005). 10.1128/jb.187.10.3502-3510.2005

[27] Uenoyama, A., Kusumoto, A., Miyata, M. Identification of a 349-kilodalton protein (Gli349) responsible for cytadherence and glass binding during gliding of Mycoplasma mobile. J Bacteriol 186, 1537–1545 (2004). 10.1128%2FJB.186.5.1537-1545.2004

[28] Aluotto, B.B., Wittler, R.G., Williams, C.O., Faber, J.E. Standardized bacteriologic techniques for the characterization of Mycoplasma species. Int. J. Syst. Bacteriol. 20, 35–58 (1970). 10.1099/00207713-20-1-35

[29] Miyata, M., Yamamoto, H., Shimizu, T., Uenoyama, A., Citti, C., Rosengarten, R. Gliding mutants of Mycoplasma mobile: relationships between motility and cell morphology, cell adhesion and microcolony formation. Microbiology 146, 1311–1320 (2000). 10.1099/00221287-146-6-1311

[30] Nakane, D., Miyata, M. Mycoplasma mobile cells elongated by detergent and their pivoting movements in gliding. J Bacteriol 194, 122–130 (2012). 10.1128%2FJB.05857-11

[31] Miyata, M., Uenoyama, A. Movement on the cell surface of the gliding bacterium, Mycoplasma mobile, is limited to its head-like structure. FEMS Microbiol Lett 215, 285–289 (2002). 10.1111/j.1574-6968.2002.tb11404.x

[32] Jaffe, J.D., Stange-Thomann, N., Smith, C., DeCaprio, D., Fisher, S., Butler, J., et al. The complete genome and proteome of Mycoplasma mobile. Genome Res 14, 1447–1461 (2004). 10.1101%2Fgr.2674004

